# The Influence of Lymphoid Reconstitution Kinetics on Clinical Outcomes in Allogeneic Stem Cell Transplantation

**DOI:** 10.1101/065680

**Authors:** David J Kobulnicky, Roy T Sabo, Shashank Sharma, Ali S Shubar Ali, Kristen Kobulnicky, Catherine H Roberts, William B Clark, Harold Chung, John M McCarty, Amir A Toor

## Abstract

Lymphoid recovery following myeloablative stem cell transplantation (SCT) displays a logistic pattern of exponential growth followed by a plateau. Within this logistic framework, lymphoid recovery is characterized by the parameters *R* (slope of ascent), *a* (time of maximal rate of ascent) and *K* (plateau), the steady state lymphocyte count. A retrospective analysis of allogeneic SCT performed from 2008 to 2013 was undertaken to compare lymphoid recovery and clinical outcomes in 131 patients with acute myelogenous leukemia, acute lymphocytic leukemia and myelodysplastic syndromes. Using *Prism* software, a logistic curve was successfully fit to the absolute lymphocyte count recovery in all patients. Patients were classified according to the magnitude and rate of lymphoid recovery; pattern A achieved an ALC of > 1000/µL by day 45; pattern B an ALC 500 < *x* < 1000 /µL and pattern C, an ALC <500/µL. Pattern A was characterized by a higher mean K (p < 0.0001) compared with patterns B & C. Patients with patterns B and C were more likely to have mixed T cell chimerism at ninety days following SCT (p=0.01). There was a trend towards improved survival, (and relapse-free survival) in those with pattern A and B at one year compared to pattern C (p = 0.073). There was no difference in cGVHD (p=0.42) or relapse (p=0.45) between pattern types. CMV, aGVHD and relapse all were heralded by deviation from logistic behavior. Pattern C patients were more likely to require donor lymphocyte infusion (p=0.017). Weaning of Tacrolimus post-transplant was associated with a second, separate logistic expansion in some patients. This study demonstrated that lymphoid reconstitution follows a prototypical logistic recovery and that pattern observed correlates with T cell chimerism and need for DLI, and may influence survival.

## INTRODUCTION

Predicting the clinical trajectory of any stem cell transplant (SCT) recipient is difficult given the many factors that influence the likelihood of relapse or graft verse host disease (GVHD) in each individual. Current prediction models rely upon statistics and probability theory to estimate the likelihood that an event will occur in a population at a given moment but cannot inform as to the trajectory of an individual patient.^1^ Static or discrete measures, such as lymphocyte counts at specific times following SCT, conceptualize outcomes in terms of discrete functions in a population of patients and therefore only yield probability distributions for clinical outcomes in individual patients.^2^ A prognostic model that captures data as a continuous function of time, may on the other hand, account for the evolution in clinical parameters and may more successfully correlate them with clinical outcomes in unique individuals.

Biological events, such as GVHD or relapse that occur in the course of transplantation are governed by immune reconstitution following SCT.^3^ Immune reconstitution is a dynamic process occurring over time following SCT. As such, bone marrow transplant may be considered as a dynamical system, much like a growing population, in which lymphoid recovery in the early days following SCT may dictate the trajectory of future outcomes.^4^, ^5^ Therefore, quantitative measures of lymphocyte recovery over time may be able to predict outcomes and help guide therapies.^6^, A newly recognized feature of lymphoid reconstitution is that it is a logistic function of time. This is not surprising given that the logistic curve is ubiquitously observed when measuring growth in biological systems. ^7^ In such systems, exponential growth occurs until environmental resources are constrained, it then plateaus reaching a steady state. Lymphoid recovery post-transplant is similar. ^8^ Lymphoid cells, comprised of NK, T and B cell clones compete for hematopoietic resources such as cytokines, driven by pathogen as well as tumor and self-antigens and eventually the population of lymphocytes stabilizes with a dominant and minority clones present. The balance of alloreactive and pathogen directed clones may influence the clinical trajectory a patient may follow.

Systems growing as a logistic function of time are quantitatively characterized by a starting magnitude (*N_0_*), an initial period of slow growth followed by a period of exponential expansion, resulting in a steady state (*K*) population after the passage of time (*t*). The exponential period of growth is characterized by specific rate of growth (*R*), as well as the time to maximal growth, the point of maximal inflection (*a*). These variables are represented in the following equation: *N_t_= N_0_+(K-N_0_)/(1+10 ^(a-t)R)^* providing a value *N_t_* for any given period of time.^9, 10^ Previous work has suggested that a general logistic curve could be fit to lymphoid recovery post-transplant occurring following reduced intensity conditioning and SCT, and thus was chosen as the model for this study.

Absolute lymphocyte counts (ALC) are monitored frequently and regularly post-transplant and provide adequate data to allow an examination of immune recovery kinetics, and test hypotheses about relationship between logistic parameters and later clinical outcomes^11^. Admittedly, ALCs are only an indirect estimate of the risk for either GVL or GVHD but their frequency permits analysis of the growth kinetics as the subtleties of plateau, growth and inflection can be easily identified. All the parameters vary, influenced by the many biologic and pharmacologic factors post-transplant such as HLA mismatch, infection, conditioning, GVHD prophylaxis and many more. ^12, 13, 14, 15,^ Plotting ALC as a continuous function of time is useful in capturing the dynamic and evolutionary change over time and offers an alternative to static or discrete analysis.

This study aims to determine whether lymphoid recovery following myeloablative conditioning follows logistic kinetics, and whether logistic modeling can be used to predict outcomes and thus guide therapies to minimize the competing pathologies of GVHD and relapse. Data for this study originate from a retrospective analysis of patients transplanted from 2004-2011 that compared outcomes between HLA matched related donor (MRD) SCT recipients, with HLA matched unrelated donor (MUD) recipients who received ATG for GVHD prophylaxis.^16, 17, 18, 19, 20^ That study found comparable survival, relapse and immune reconstitution between MRD and MUD recipients where the latter received ATG peri-transplant. ^21^ It was observed that patient’s with earlier and stronger lymphoid recovery appeared to have improved survival and freedom from relapse but perhaps more GVHD.

## MATERIALS and METHODS

### Patients

After obtaining permission from the Virginia Commonwealth University Institutional Review Board, and in accordance with the declaration of Helsinki, we conducted a retrospective review of the medical records for allogeneic SCT recipients with AML, ALL, MF or MDS transplanted between 2008 and 2013. Patients were followed for a minimum of one year and for as long as existing records permitted. To be eligible for inclusion, patients had to be older than 18 years of age, previously diagnosed with acute myelogenous leukemia, acute lymphocytic leukemia or myelodysplastic syndrome and have undergone myeloablative allogeneic SCT from a MUD or MRD ^22^. All patients were matched at 8/8 or 7/8 HLA-A,-B,-C and-DRB1 loci. In general, MUD underwent high resolution typing, while MRD received intermediate resolution typing. Conditioning regimens included total body irradiation (TBI) (12 Gray, given over days -6 to -4) or Busulfan (0.8mg/kg IV every 6 h, days -7 to -3) given along with cyclophosphamide (60mg/kg/day IV, days -3 and -2); alternatively Busulfan (130mg/m^2^/day IV days -5 to -2) with Fludarabine (40mg/m^2^/day IV days -5 to -2), or Fludarabine (30mg/m^2^/day IV days -6 to -3) with Melphalan (140 mg/m^2^/day IV day -2) were administered. Patients who received an unrelated SCT received 5 mg/kg rabbit Anti-Thymocyte Globulin (ATG) in divided doses from days -3 to -1 (Thymoglobulin, Sanofi-Aventis, Quebec, CA). Patients who received a MRD transplant typically received Cyclosporine and Methotrexate as GVHD prophylaxis while patients who received a MUD typically received Tacrolimus and Methotrexate.

### Study Design

Absolute lymphocyte counts (µL^-^1) from the day of transplant until 180 days afterwards were extracted from the medical record and recorded. The ALC values were graphed over time following SCT and inspected for the presence of logistic growth. Subsequently, using Graph Pad Prism version 6.0 (GraphPad Software, La Jolla, California), a logistic function was fit to the lymphoid recovery curves over time for days 0 to 60 or until lymphoid recovery deviated from the logistic. A logistic function defined as *N_t_= N_0_+(K-N_0_)/(1+10 ^(a-t)R)^* was applied to individual patient’s lymphoid recovery. The quantitative parameters of *N_0_*, *K*, *R*, and *a* were recorded for each patient. *N_0_* represents the lymphocyte count at the time of transplant, *K* represents the lymphocyte count at ‘steady state’ which is usually the peak value. *N_t_* represents the lymphocyte count at time *t* following transplant, *a* represents the time at which growth rate is maximal and at which an inflection point is observed in the logistic curve. *R* represents the rate of growth and is estimated by the slope of the logistic curve. The clinical event that caused deviation from the logistic was noted for all patients in whom there was a deviation recorded from the logistic behavior. For patients in whom a clinical event occurred prior to achievement of *K*, the peak absolute lymphocyte prior to the clinical event was used as the ‘steady state’ value. Clinical events including relapse, aGVHD, cGVHD and infection as well as their attendant therapies were catalogued as well to determine if they were temporally related to deviation from the generally observed logistic pattern. Donor engraftment was measured using chimerism analyses performed at approximately 12, weeks following transplant on whole blood, bone marrow and immunomagnetically separated total T cells. Acute (aGVHD) and chronic GVHD (cGVHD) were also assessed. All patients had their aGVHD scored with the IBMTR grading scale ^23^, ^24^ and cGVHD scored with the Seattle grading system. ^25^ For purposes of analysis, late-onset aGVHD symptomatology was categorized as aGVHD and ‘overlap syndrome’ symptomatology as cGVHD.

### Statistical Analysis

The primary study end points were survival, relapse-free survival, and GVHD. Survival and relapse-free survival (any event of fatality, relapse) for different logistic patterns (designated, patterns A, B and C) were plotted with Kaplan-Meier curves and compared using the log-rank test. The incidence curves for relapse, acute GVHD, chronic GVHD, and DLI – accounting for the competing risk of fatality – are also plotted with step-wise curves, and compared between groups using Grey’s test. The Cox proportional hazards model is used to model time-to-event outcomes against the continuous logistic parameters (*K* and *R*). Categorical patient identifiers (diagnosis, donor pattern, stem cell source, race, gender, and conditioning regimen) were compared between patients with differing lymphoid recovery patterns using chi-square or exact tests. Numerical patient measures (*a*, *K* and *R*) are compared between patterns using analysis of variance, where reported p-values and confidence intervals of differences between means are adjusted for multiple comparisons using the Tukey approach. The *survival* package in the R statistical software (version 2.15) was used for all time-to-event analyses. The SAS statistical software (version 9.4) is used for all other analyses, with the PHREG procedure used to fit Cox proportional hazard models, and the FREQ procedure used for chi-square and exact tests.

## RESULTS

### Patient Characteristics

Between 2008 and 2013, one hundred and thirty-one patients underwent allografting at Virginia Commonwealth University (Table 1). Fifty-six patients were MRD recipients and received no ATG as a part of their conditioning, while seventy-five were MUD recipients and received ATG as part of conditioning. There was no difference in diagnosis, conditioning regimen, race, age, CMV susceptibility, mean CD3 or CD34 cell dose infused. MRD were significantly more likely to have received cyclosporine for GVHD prophylaxis (32/56) while MUD were more likely to Tacrolimus GVHD prophylaxis (65/76, p=<0.0001). MUD were more likely to be HLA mismatched (17/76) than MRD (1/55, p=0.011).

**TABLE 1.** Patient characteristics by ALC recovery kinetic pattern. *1: TBI / Etopside, Fludarabine / Melphalan; *2: No ATG used in conditioning, CyclosporinàTacrolimus; *3: Rabbit-ATG used in conditioning.*4: Tacrolimus à Corticosteroids, Tacrolimus à Cyclosporine, Tacrolimus + MMF

|  | MRD | MUD | p-value |
| --- | --- | --- | --- |
| *N* | 56 | 75 |  |
| *Gender* (Female) | 28 | 34 | 0.60 |
| *Race* (Caucasian) | 45 | 60 | 0.99 |
| *Age* (years: median) | 52.7 | 49.1 | 0.14 |
| *Conditioning Regimen* |  |  | 0.96 |
| Busulfan / Cyclophosphamide | 16 (29%) | 21 (28%) |  |
| TBI / Cyclophosphamide | 19 (34%) | 24 (32%) |  |
| Busulfan / Fludarabine | 18 (32%) | 27 (36%) |  |
| Other<sup>*1</sup> | 3 (5%) | 3 (1%) |  |
| *Stem cell source* |  |  | <0.0001 |
| Bone Marrow | 0 | 14 |  |
| Peripheral Blood | 56 | 61 |  |
| *Diagnosis* |  |  | 0.97 |
| Myelodysplasia | 16 | 22 |  |
| Acute Myeloid Leukemia | 30 | 40 |  |
| Acute Lymphocytic Leukemia | 10 | 12 |  |
| *HLA Mismatch* | 1 | 17 | <0.0001 |
| *GVHD Prophylaxis* |  |  | <0.0001 |
| Cyclosporin | 32 | 2 |  |
| Tacrolimus | 10 | 65 |  |
| Other<sup>*2</sup> | 14 | 8 |  |
| CMV Susceptible | 35/55 | 45/73 | 0.38 |
| ABO Incompatibility | 16 | 22 | 0.54 |
| *GVHD Prophylaxis<sup>\*4</sup>* |  |  |  |
| Cyclosporin | 35 | 35 | 15 |
| Tacrolimus | 30 | 15 | 1 |
| *Cell Dose* |  |  | 0.34 |
| CD3 (10<sup>6</sup> cells/kg) | 3.35 | 2.81 |  |
| CD34 (10<sup>8</sup> cells/kg) | 4.53 | 4.20 |  |

\*1: TBI / Etoposide, Fludarabine / Melphalan; \*2: No ATG used in conditioning, Cyclosporin → Tacrolimus; \*3: Rabbit-ATG used in conditioning.
\*4: Tacrolimus → Corticosteroids, Tacrolimus → Cyclosporine, Tacrolimus + MMF

### Lymphoid Reconstitution Kinetics

Patients were classified according to the rate and magnitude of their lymphoid recovery. Pattern A was a high and rapid lymphoid recovery defined as an ALC of 1000±100 µL^-1^ for three consecutive days by day 45 (Figure 1). Pattern B was a lower and slower lymphoid recovery that did not meet the criteria for Pattern A, but achieved an absolute lymphocyte count of at least 500±100 µ^L-^1 for three consecutive days by day 45. Pattern C was a delayed and inadequate lymphoid reconstitution and inclusive of all patients who did not meet criteria for either pattern A or B. Patient characteristics were then analyzed according to lymphoid recovery pattern they experienced. Day 45 was chosen because nearly all patients had experienced a clinical plateau in lymphocyte count by this time. ALC at this time point therefore was determined to be a good surrogate for *K*. Patients with pattern A had significantly larger mean *K* than either pattern B (p < 0.0001) or C (p < 0.0001) but Group B and C were not significantly different (p = 0.0826, Table 3). Neither mean *a* (p = 0.3657, Table S2), nor *R* (p = 0.1439, Table S3) were significantly different between pattern different patient groups. Mean *a* was later in MUD than MRD recipients (*u*-MRD=19.0, *u*-MUD=23.8 days post transplant, p=0.32). There was no difference in conditioning regimen, stem cell source, race, age, CMV susceptibility, mean CD3 or CD34 cell dose infused between patients with different pattern types (Table 3). Leukemia was more common among patients with pattern C (p=0.031).

**FIGURE 1:**
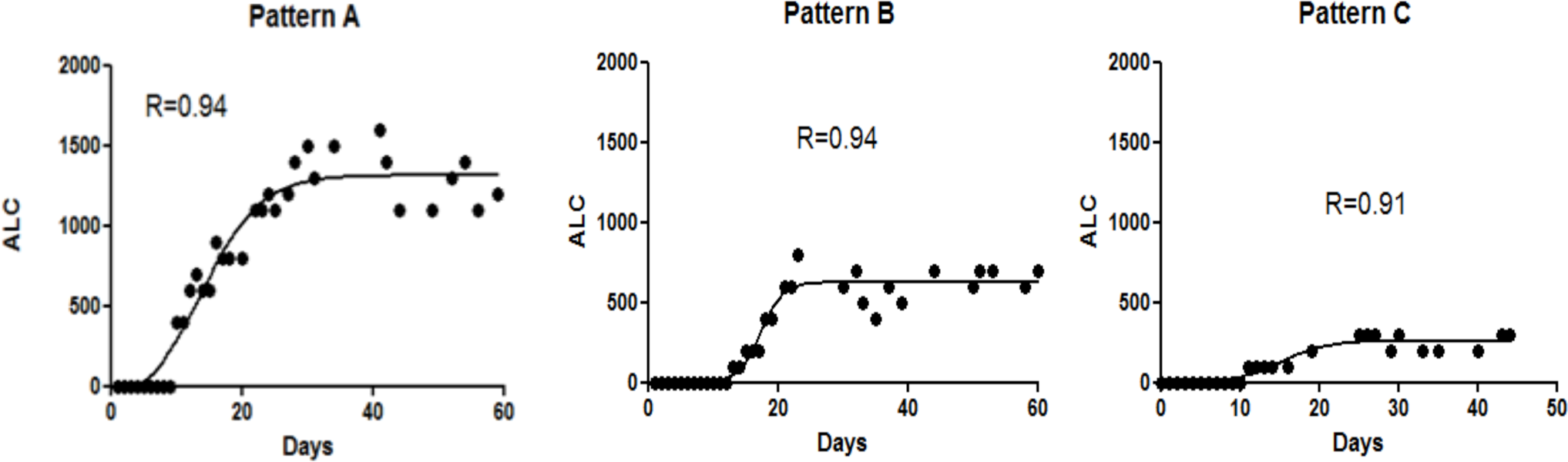
Patterns of ALC recovery. This figure demonstrates patients with prototypical pattern A, B and C lymphoid reconstruction each plot fitted with a logistic curve using Prism software

**TABLE 2.** Patient characteristics by ALC recovery kinetic pattern. *1: TBI / Etopside, Fludarabine / Melphalan; *2: No ATG used in conditioning;, CyclosporinàTacrolimus *3: Rabbit-ATG used in conditioning.*4: Tacrolimus à Corticosteroids, Tacrolimus à Cyclosporine, Tacrolimus + MMF

|  | **A** | **B** | **C** | **p-value** |
| --- | --- | --- | --- | --- |
| *N*= | 65 | 50 | 16 |  |
| Gender (Male) | 30 | 27 | 12 | 0.11 |
| Race (Caucasian) | 50 | 42 | 13 | 0.61 |
| Age (years: median) | 51.2 | 49.1 | 53.1 | 0.54 |
| *Conditioning Regimen* |  |  |  | 0.25 |
| Busulfan / Cyclophosphamide | 20 (31%) | 16 (32%) | 1 (7%) |  |
| TBI / Cyclophosphamide | 24 (40%) | 13 (26%) | 6 (37%) |  |
| Busulfan / Fludarabine | 20 (31%) | 18 (36%) | 7 (44%) |  |
| Other<sup>*1</sup> | 1 (2%) | 3 (6%) | 2 (12%) |  |
| *Stem Cell Source* |  |  |  | 0.10 |
| Bone Marrow | 7 | 3 | 4 |  |
| Peripheral Blood | 58 | 47 | 12 |  |
| *Donor Relation* |  |  |  | **0.014** |
| Related<sup>*2</sup> | 34 | 20 | 2 |  |
| Unrelated<sup>*3</sup> | 31 | 30 | 14 |  |
| *Diagnosis* |  |  |  | 0.031 |
| Myelodysplasia | 20 (31%) | 18 (36%) | 0 (0%) |  |
| Acute Myeloid Leukemia | 35 (54%) | 25 (50%) | 11 (69%) |  |
| Acute Lymphocytic Leukemia | 10 (15%) | 7 (14%) | 5 (31%) |  |
| HLA Mismatch | 5 | 10 | 3 | **0.011** |
| *GVHD Prophylaxis<sup>\*4</sup>* |  |  |  | **0.0071** |
| Cyclosporin | 22 | 11 | 0 |  |
| Tacrolimus | 33 | 27 | 14 |  |
| Other<sup>*2</sup> | 10 | 12 | 2 |  |
| CMV Susceptible | 45 | 24 | 12 | **0.03** |
| ABO Incompatibility | 19 | 14 | 5 | 0.96 |
| *Cell Dose* |  |  |  |  |
| CD3 (10<sup>6</sup> cells/kg) | 3.43 | 2.96 | 1.94 |  |
| CD34 (10<sup>8</sup> cells/kg) | 4.58 | 4.48 | 4.18 |  |

\*1: TBI / Etoposide, Fludarabine / Melphalan; \*2: No ATG used in conditioning; Cyclosporin → Tacrolimus \*3: Rabbit-ATG used in conditioning.
\*4: Tacrolimus → Corticosteroids, Tacrolimus → Cyclosporine, Tacrolimus + MMF

**TABLE 3.** Logistic parameter K and Pattern Type. Steady state ALC values (L^-^1) in patients demonstrating different ALC recovery kinetics. N=number, SE= Standard of Error, CI = Confidence Interval DIFF : How groups are compared. “A-B” signifies that pattern A is being compared against pattern B, and so on.

|  | **K** |  |  |  |  |  |  |
| --- | --- | --- | --- | --- | --- | --- | --- |
| Group | N | Mean | SE | 95% CI | Diff | CI | p-value |
| A | 65 | 1765.2 | 133.3 | 1501.3, 2029.1 | A-B | 415.8, 1220.8 | < 0.0001 |
| B | 50 | 946.8 | 153.6 | 642.9, 1250.8 | A-C | 755.7, 2092.7 | < 0.0001 |
| C | 16 | 340.9 | 310.3 | -273.3, 955.1 | B-C | -79.4, 1291.2 | 0.0826 |

### Pattern Type and Donor Engraftment

Lymphoid reconstitution kinetics in the first month following SCT were predictive of donor engraftment, as pattern type was subsequently able to influence the incidence of mixed chimerisms at 90 days post transplant. Mixed T cell chimerism (p = 0.0139, Table 4) was significantly more likely in those with pattern C than patterns A or B, and more likely in pattern B than pattern A. Likewise, mixed whole blood chimerism was significantly more likely in those with pattern C lymphod reconstitution (p =0.0414, Table 5). Similarly, there is a trend towards mixed bone marrow chimerisms in pattern C patients (p = 0.0996). Few patients who developed mixed T cell chimerisms early post transplant later achieved fully donor states but no events were seen at all in patients with poor reconstitution (Pattern A: 2/13, Pattern B: 3/16, Pattern C: 0/9).

**TABLE 4.** T Cell Chimerism at 90 days following SCT and lymphoid reconstitution pattern type.

| Logistic Group | At least 95% T-Cell Chimerism |  |  |
| --- | --- | --- | --- |
|  | Yes | No | Total |
| A | 52 (80%) | 13 (12%) | 65 |
| B | 34 (68%) | 16 (32%) | 50 |
| C | 7 (44%) | 9 (56%) | 16 |

**TABLE 5.** Whole Blood Chimerism at day 90 and Pattern Type.

| Logistic Group | At least 95% Whole Blood Chimerism |  |  |
| --- | --- | --- | --- |
|  | Yes | No | Total |
| A | 48 (76%) | 15 (24%) | 63 |
| B | 35 (70%) | 15 (30%) | 50 |
| C | 7 (44%) | 9 (56%) | 16 |

### Clinical Outcomes

In these recipients of myeloablative conditioning and allografts for leukemia and MDS, the chimerism trends in patients with different lymphoid reconstitution kinetics were reflected in the clinical outcomes. There is a trend towards improved survival in patterns A and B compared to C (chi-square p =0.073, Figure 2). 78% of patients with pattern A were alive at one year compared to only 66% of those with Pattern B and 37.5% with Pattern C. Similar trends were observed for relapse (chi-square p =0.4507, Figure S1) and relapse-free survival (RFS, chi-square p = 0.1360, Figure S2); 65% of patients with Pattern A had not relapsed by on year as compared to only 58% of those with pattern B and 38% of pattern C. Patients with pattern C often experienced early death due to infection or peri-transplant complication and making true relapse incidence estimates difficult.

**FIGURE 2:**
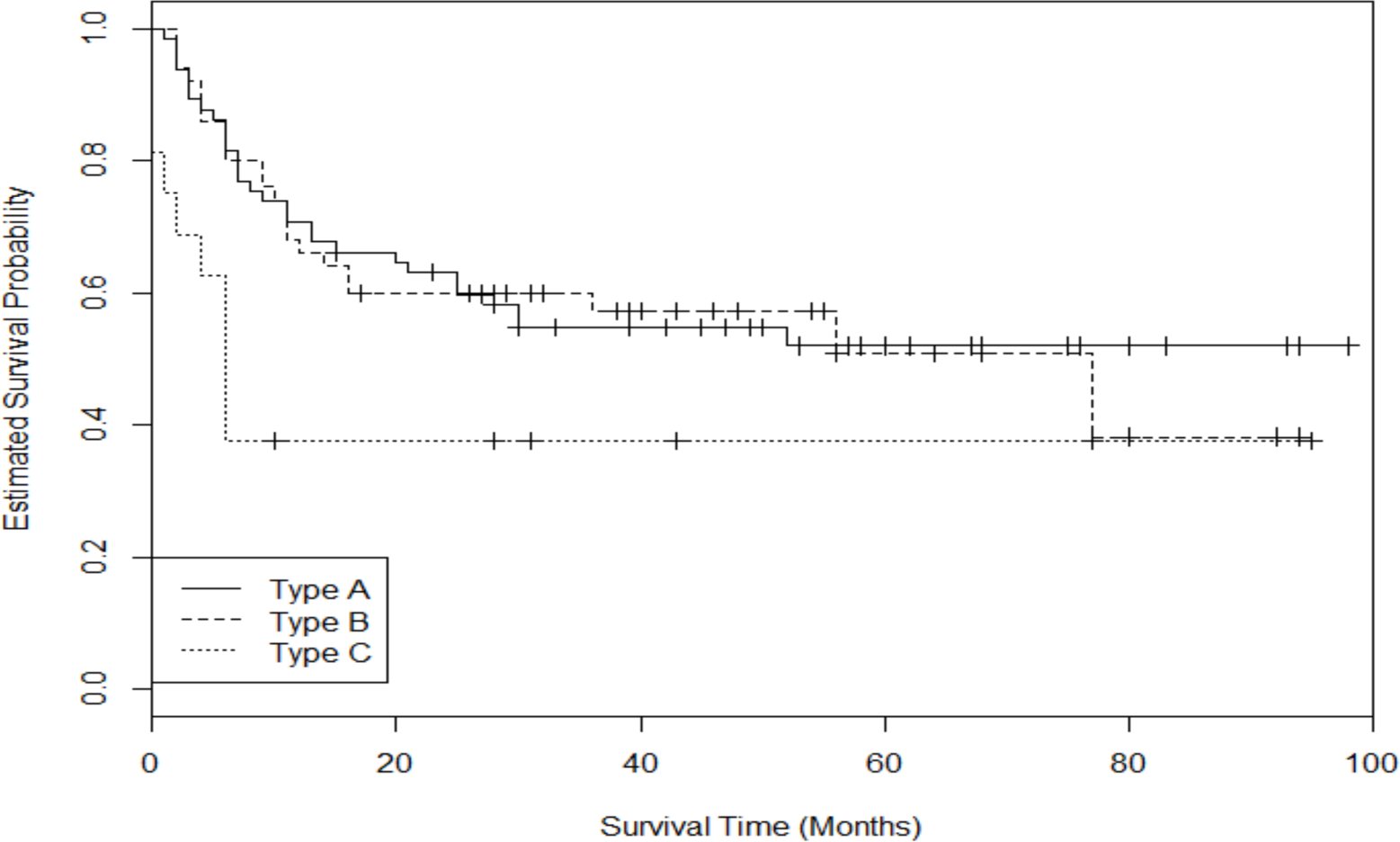
Kaplan-Meier survival plot of patients grouped according to ALC recovery pattern.

Acute GVHD (Figure 3) while not statistically significantly different between pattern types (chi-square p = 0.2068) was higher in patients with patterns A and B. Only 31% of patients with Pattern C experienced aGVHD as compared to 57% of those with pattern A and 62% with pattern B. Chronic GVHD (Figure S3) was not significantly different between pattern type (chi-square p = 0.4183). However, MUD recipients (23/31=81%) with pattern A were significantly more likely to experience cGVHD than MRD recipients (12/34=41%, p = 0.0012). There were no differences in acute GVHD, survival, relapse, or relapse-free survival when pattern type was segregated by donor type.

**FIGURE 3:**
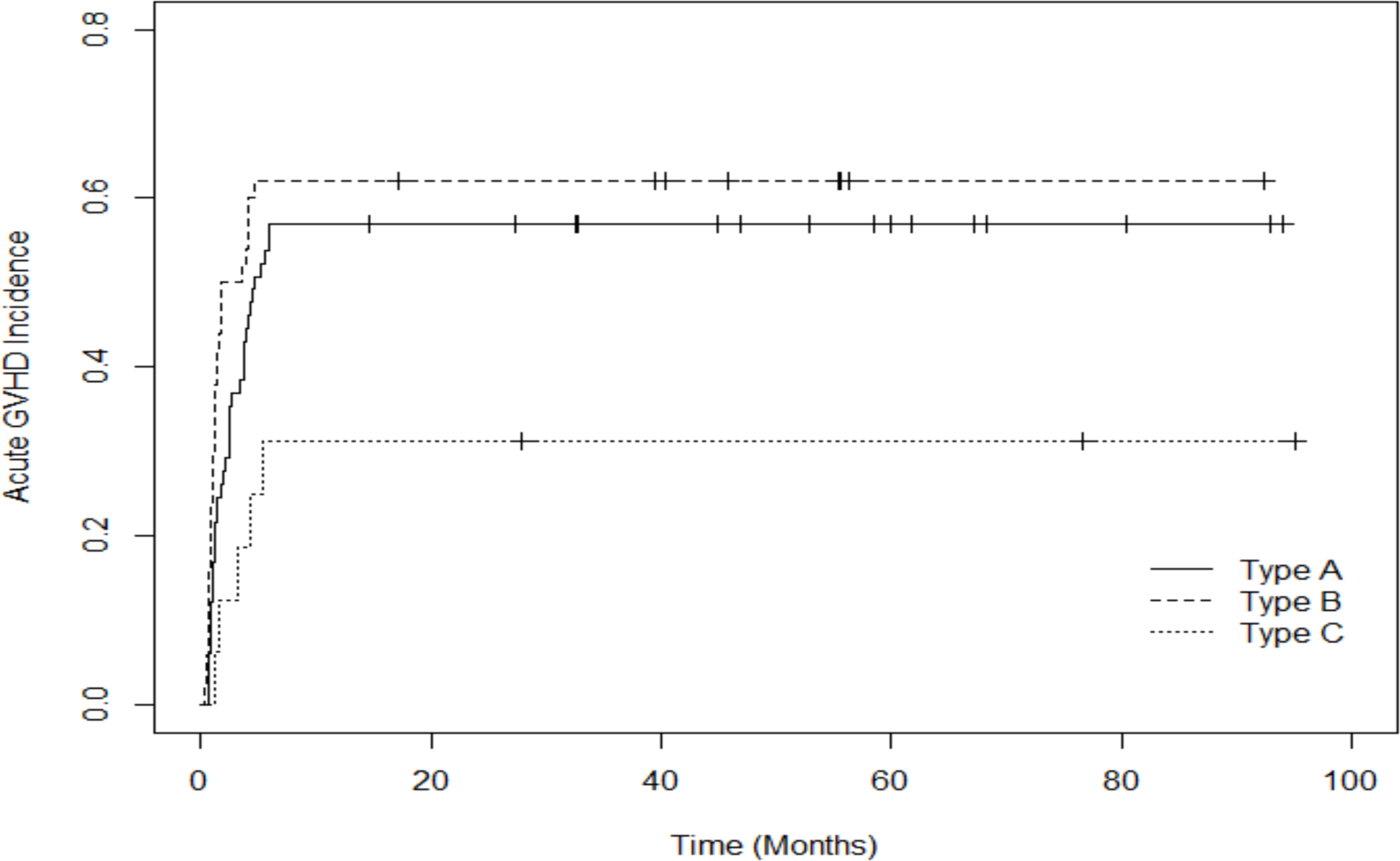
Cumulative Incidence of Acute GVHD plot of patients according to ALC recovery pattern.

Finally, patients with poor reconstitution were more likely to require donor lymphocyte infusion (DLI) than those with avid reconstitution (Figure 4, p=0.0170). Only 7% of patients with Pattern A required DLI compared to 18% with Pattern B and 18.8% with Pattern C.

**FIGURE 4:**
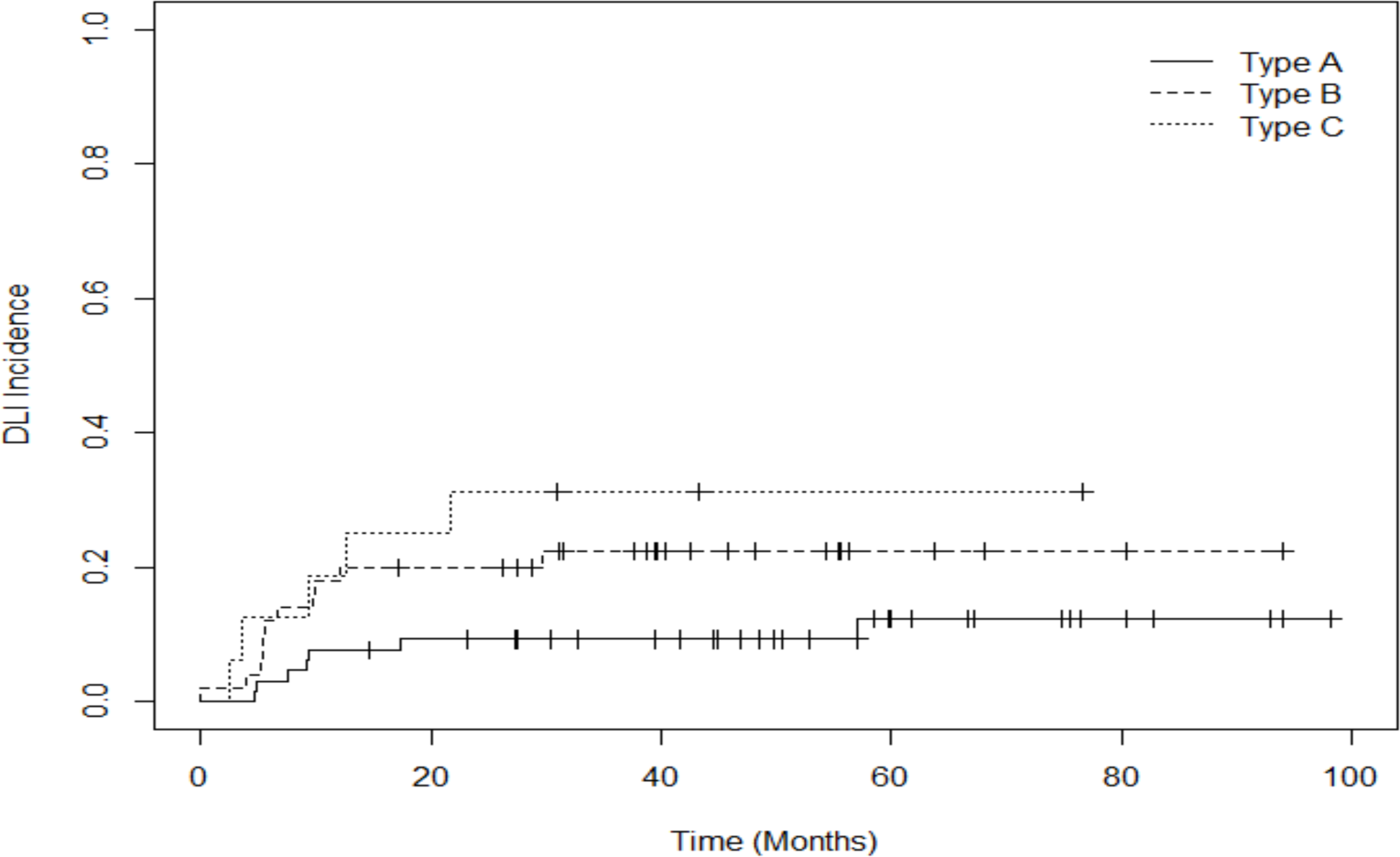
Need for DLI over time.

### Phenotypes and Portraits of Immune Reconstitution

As lymphoid recovery over time was recorded and clinical events mapped alongside, many interesting observations emerged. Many patients experienced an early peak in their lymphocyte counts that later stabilized (Figure 5). Many had a more significant pre-plateau peak in their lymphocyte count often more than 1,000 µL^-1^ higher or 25% greater than the following plateau. In nearly all cases, we were able to find an associated clinical event, usually aGVHD (A: 38/65, B: 31/50, C: 5/16, Figure 6) or infection (A: 40/65, B: 18/26, C: 14/16). Within the overall larger logistic trend, there were often periods demonstrating minor logistic growth indicating periods of lymphoid expansion resulting in a new higher ALC plateau. Two notable examples were, Tacrolimus withdrawal near day 100 (Figure 5) and the cessation of MMF at Day 42 (Figure S4). This makes intuitive sense; in population ecology language, the carrying capacity (maximum population, K) permitted by the environment is increased as immunosuppression is withdrawn and the T cell growth rate increases. As therapies were instituted, such as, antibiotics or steroids, some patients’ ALC curves trended back towards steady state, logistic behavior. Relapse or GVHD therapy marked a dramatic, negative deflection from this plateau and was rarely subtle (A:19/65, B:12/50, C: 5/18, Figure S5). Alternately, viral infection and GHVD onset was usually a positive deflection. Examples of eighty-two patients’ lymphoid reconstitution are included categorized by Pattern Type (Figure, S6). Side by side comparison reveals both the similarities and contrasts between patients but also highlights the importance of the trend as opposed to a value in isolation.

**FIGURE 5:**
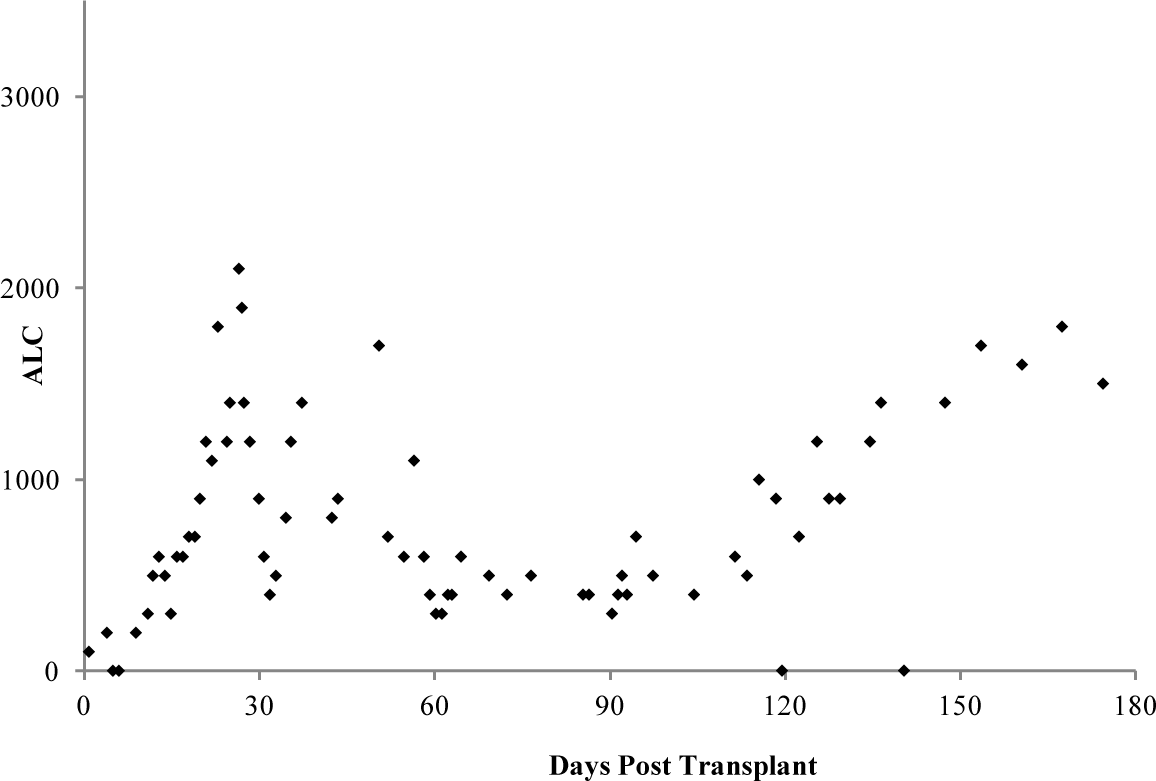
Example of early peak in otherwise normotypic logistic behavior. This early peak at day 20 was common in many patients. This patient peaks to an ALC of 2200/? L with eventual plateau 700/? L – this was not a significant deviation from logistic behavior and clinical significant aGVHD or infection could not be detected. Again, as tacrolimus is weaned, a second later logistic rise occurs with a new plateau around day 150.

**FIGURE 6:**
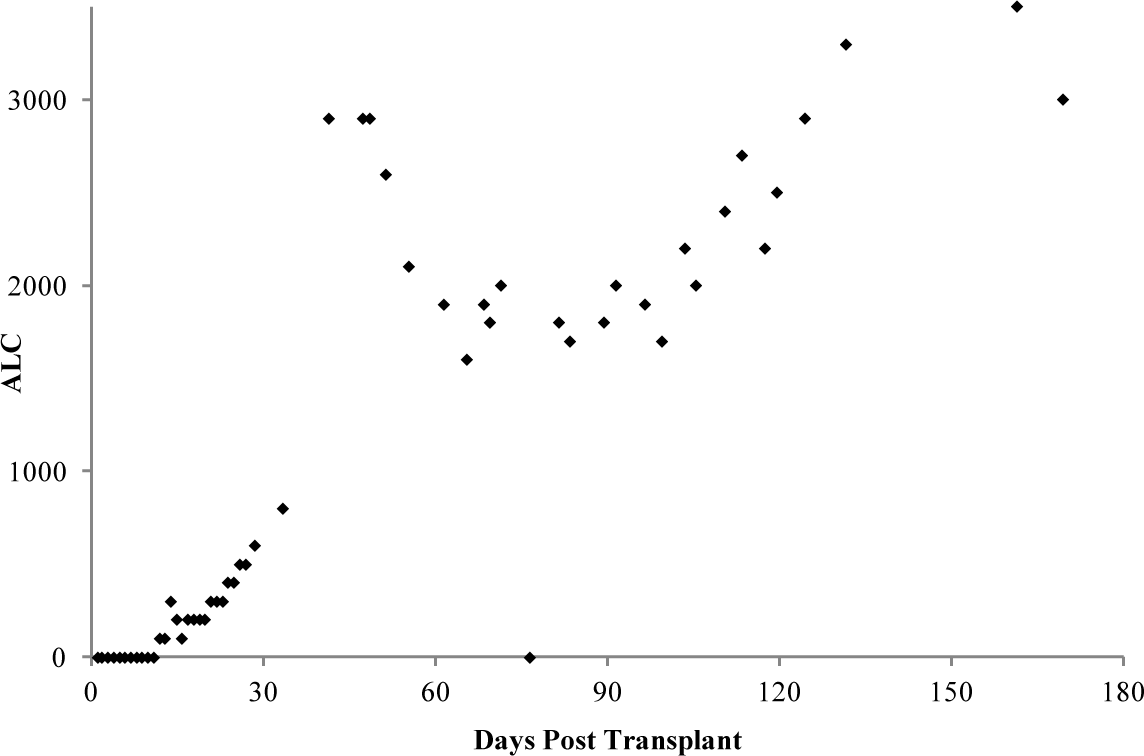
Tacrolimus and aGVHD influencing logistic behavior. This patient experiences grade III aGVHD around Day 40 and has a peak around ~ 3000 /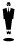 L. As their aGVHD is treated, the eventual plateau resets around ~ 2000/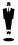 L from Day 60 to 90. As tacrolimus is weaned, a second, slower logistic rise can be observed from Day 100 to Day 140.

## DISCUSSION

Immune reconstitution is critical to successful stem cell transplantation with a number of studies demonstrating the salutary effect of early lymphocyte recovery on clinical outcomes. Most of these studies however report association of clinical outcomes with lymphoid reconstitution at specific times following SC. This method of analysis, while adequate for prognostication, does not contribute to developing a comprehensive understanding of the biology of post transplant immune reconstitution. In this study we report the lymphocyte reconstitution kinetics in a large cohort of patients undergoing allogeneic SCT and demonstrate that they follow a logistic growth pattern. This implies that post transplant immune reconstitution may be studied in a quantitative fashion over time, and a deeper understanding of its relationship of lymphoid recovery kinetics with clinical outcomes may be established. The key finding is that nearly all patients demonstrate a period of exponential expansion around 2.5 to 3 weeks following SCT followed by a plateau, the magnitude of which correlates with later T cell chimerism and eventual clinical outcomes.

Lymphocyte subsets include unique T cell clones responding to their cognate antigens. An initial upsurge is likely a consequence of inflammation causing a surge of growth, with later decline to a steady state level. This means that the early peak observed is likely related to a multitude of T cell clones growing in response to their target antigens, each growing rapidly and competing for resources. As dominant clones emerge, the population stabilizes. The clones that emerge may select for the clinical trajectory to come – GVL, GVHD or some salutory balance between the two. Lymphocyte subset analysis and T cell sequencing might illimunate the details of this cloncal competition. Presumably an oligoclonal T cell repertoire at this point may represent a higher risk of GVHD, whilst a more polyclonal repertoire will represent normal immune reconstitution. It is also important to recognize that the exponential expansion early in the course of transplant means that the eventual trajectory of the alloreactivity to be experienced by the patient is set up early following transplant. This is supported by the patients with pattern A and B having a greater likelihood of complete donor T cell chimerism and a trend towards improved survival and relapse free survival. It is likely that in larger, more uniform cohorts of patients, lymphoid reconstitution patterns and clinical impact thereof may be more informative. Another important confounder for our analysis could be the cutt off chosen to define different patterns, these were adapted from an earlier RIC regimen study of transplant for indolent hematological malignancies, where immune reconstitution is likely more critical to outcome. In our cohort of patients getting ablative allografts for leukemia and MDS this set of values may not accurately capture the threshold for optimal immune mediated disease control. In our data set pattern C (low ALC) is a very poor prognostic sign. However, what is less clear is whether a subtle difference exists between patients with modestly low ALCs, “normal” ALCs and leukocytosis. Nevertheless declining outcomes from patterns A to B to C does support the possibility that an immune reconstitition threshold exists below which the likleihood of poor clinical outcomes is higher.

An important observation here was that in recipients of MUD SCT, either due to graft related factors (HLA mismatch, use of bone marrow) or pharmacological immunosuppression (ATG and tacrolimus) there was a higher number of patients with pattern C with generally poor outcomes. It is also noteworthy that while their was not a statistically significant difference observed, the pattern observed correlated with CD34 cell dose infused, and when considering that donor T cell chimerism at 90 days appears to be influenced by the logistic pattern recorded earlier, suggests that the logistic expansion represents a robust donor derived T cell recovery, as has been demonstrated previously. (refer our BBMT paper from last year) This knowledge potentially acquired early on after SCT will allow physicians to tailor immunosuppression or use donor lymphocyte infusion earlier to optimize clinical outcomes.

A recent study dichotomized patients by attainment of ALC of 200 µL^-1^ and showed inferior suvival and relapse free survival in those with lymphoid reconstiutution below this threshold. While their findings were generally in agreement with ours, their paradigm was not. ^26^ ^27^Individualized therapies post transplant will require a dynamic understanding of the individual patient’s clinical trajectory. A stochastic model of immune reconstitution that selects uniform values and dates from which to make clinical decisions, in isolation from the trend, will be not useful in individualizing therapies. Another study found a relationship between low ALC, poor survival and need for additional therapies (corticosteroids). ^28^

We know from a previous randomized phase II trial of non-myeloablative ATG-TBI recipients that the logistic phenomenon is observed and that Pattern (A, B or C) and K are good predictors of later clinical outcomes. How this model would fare amongst myeloablative recipients with a more diverse set of conditioning regimens, pre-treatment, diagnoses and ATG exposure was not known. Patients with borderline B-C pattern may have a clinical course more similar to C. For these patients, a heightened suspicion of complication should be present when there is low *K* and deviation from logistic behavior with particular attention to the possibility of relapse. Earlier DLI or immunotherapy could be instituted. Despite the use of ATG and tacrolimus, MUD recipients with Pattern Type A experience considerably greater cGVHD. MUD recipients with Pattern Type B reconstitution have equivalent survival but less cGVHD. Hypothetically, these patients may have a large number of alloreactive clones that proliferate but are functionally suppressed by calcineurin inhibitors. Delaying immunosuppression withdrawal in these patients may be of benefit. On the other hand patients with pattern C are more fraught with mixed chimerism, and nonrelapse mortality. Early withdrawal of immunosuppression and consideration of DLI will benefit them. Overall, logistic analysis may help sum the numerous variables that both protect and hazard patients and thus help us choose better empiric therapies in the peritransplant period.

In conclusion, these findings indicate that post transplant lymphoid reconstitution occurs with mathematically precision, following same rules as population growth. This simple quantitative assessment provides a tool which easily identifies patients at risk for adverse outcomes, very early in the course of transplant. Assesment of lymphoid recovery kinetics in uniformly treated patients in a disease specific manner will likely yield more accurate context-specific thresholds for accurate prognostication. Extrapolation of this rule to T cell subset recovery kinetics may further enhance the value of this growth kinetic analysis in allograft recipients.

## SUPPLEMENT

### Patient Characteristics

The following patient characteristics at baseline were evaluated to ensure similarity between the two cohorts: intensity of transplant conditioning, diagnosis, age of donor, age of recipient, race, gender, prognosis, CD3, CD34 dose, ATG dose, HLA incompatibility, stem cell source, conditioning regimen and intensity, immunosuppressive regimen, CMV sero-status and ABO incompatibility.

### Patient Risk Stratification

Patient’s prognosis was classified as follows. Acute Leukemia patients were classified as CR1, CR2 or >CR2. Included within >CR2 were patients who were not able to attain remission due to primary induction failure. MDS patients were classified as standard or elevated risk. Elevated risk patients were those with RAEB I-II, MDS/MPD, treatment related MDS or secondary AML. Standard risk patients included all other sub-patterns. Patients with acute leukemia were risk stratified according to number of consolidated remissions they had experienced while patients with MDS were stratified as standard or high risk based on their disease pattern.

### Logistic Parameters and Clinical Outcomes

The group-specific survival curve (excluding Group C) is not significantly affected by either parameter *K* (p = 0.4244), *a* (p = 0.1971) or *R* (p = 0.9810). The group specific relapse curve (excluding Group C) is not significantly effected by *K* (p = 0.7760), *a* (p = 0.3784) or *R* (p = 0.9335). The group-specific acute GVHD curves (excluding Group C) are not significantly affected by K (p = 0.1396), A (p =0.8839) or R (p = 0.8284). The group-specific chronic GVHD curves (excluding Group C) are not significantly affected by *K* (p = 0.1880), *a* (p = 0.3652) or *R* (p = 0.1047). The group-specific DLI curves (excluding Group C) are not significantly affected by *K* (p = 0.6077), *a* (p = 0.7349) or *R* (p = 0.1009). There was no difference in either *K* (*u*-MRD=1350, *u*-MUD=1281, p=0.75) or *a* (*u*-MRD=0.43 *u*-MUD=0.39, p=0.78) between MRD and MUD SCT recipients.

### Clinical Outcomes by Donor Type

Within Group A patients, there was a significant difference in chronic GVHD rates between donor types (p = 0.0012), where MUD patients (23/31=81%) were more likely to experience cGVHD than MRD patients (12/34=41%). There were no significant differences in Group A between donor types 20 in rates of acute GVHD (p = 0.2378), death (p = 0.3992), relapse (p = 0.6084), DLI (p = 0.5961), event-free survival (exact p = 0.6744), or relapse-free survival (p = 0.5448). There were no significant differences in Group B between donor types in rates of acute GVHD (p = 0.8120), death (p = 0.2971), chronic GVHD (p = 0.0922), relapse (exact p = 0.9999), DLI (exact p = 0.9999), event-free survival (p =0.9999), or relapse-free survival (p = 0.2952).

**Figure S1.**
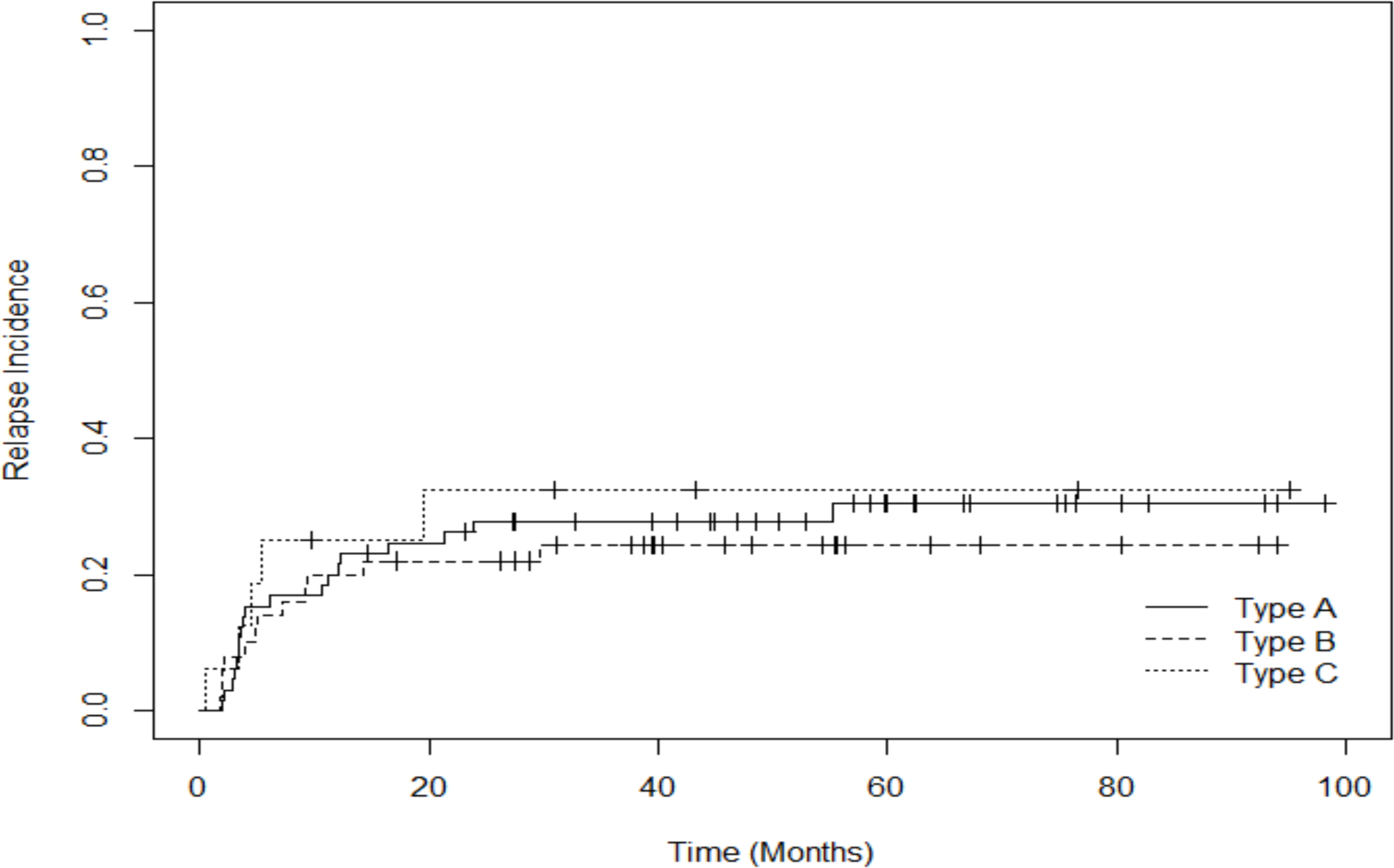
Cumulative Incidence of Relapse plot grouped according to ALC pattern.

**FIGURE S2:**
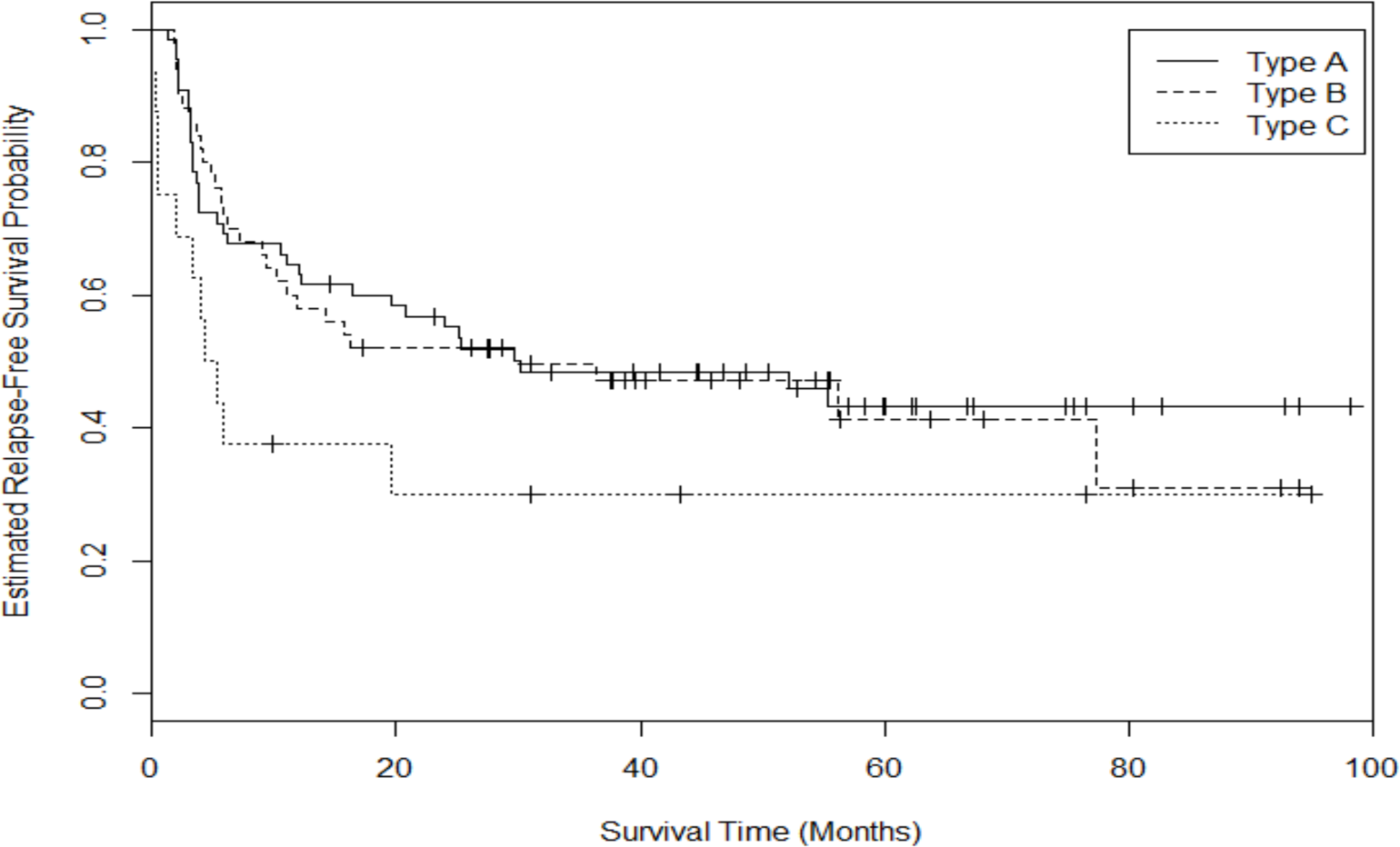
Kaplan-Meier event free survival plot of patients grouped according to pattern.

**Figure S3:**
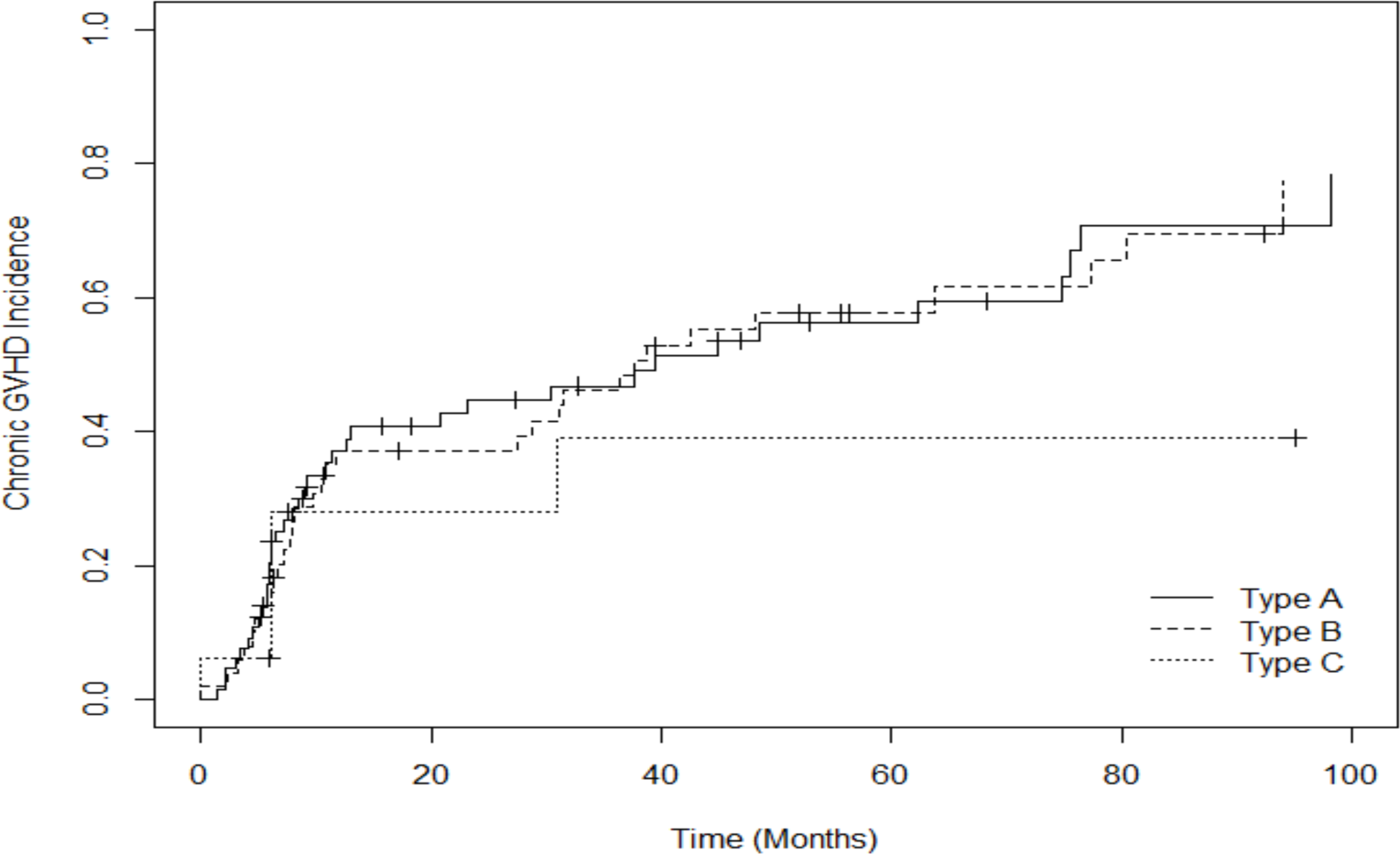
Cumulative Incident of Chronic GVHD plot according to ALC pattern.

**Figure S4:**
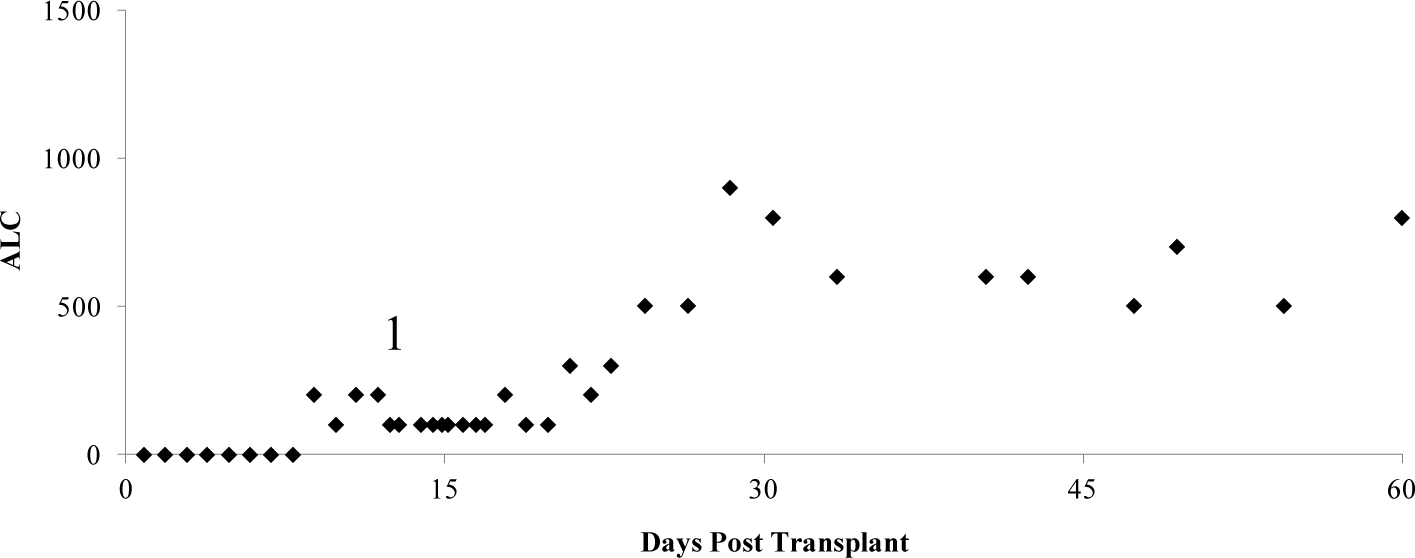
Methotrexate Influencing Logistic Behavior. When methotrexate was stopped (1) at Day 14, nearly all patients had a minor logistic rise in ALC, coincident with hematopoietic engraftment detectable from days 14 to 30.

**Figure S5:**
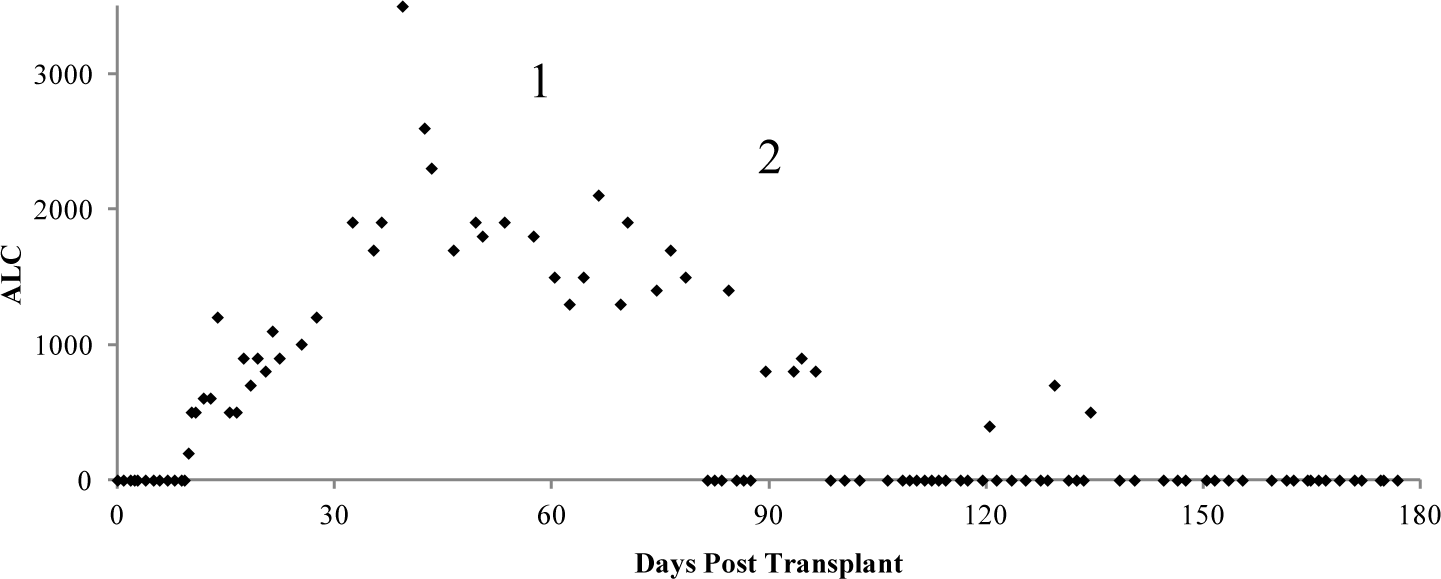
Relapse influencing logistic behavior. This patient experiences lymphoid reconstitution with Pattern A recovery and had a clinical plateau (1) in ALC from Day 40 until relapse (2) on day 75 causing an abrupt deviation from logistic behavior.

**TABLE S1.** Bone Marrow Chimerism and ALC recovery pattern

| Logistic Group | At least 95% Bone Marrow Chimerism |  |  |
| --- | --- | --- | --- |
|  | Yes | No | Total |
| A | 48 (75%) | 16 (25%) | 64 |
| B | 33 (67%) | 16 (33%) | 49 |
| C | 7 (47%) | 8 (53%) | 15 |
| Total | 88 | 40 | 128 |

**TABLE S2.** Logistic parameter a and Pattern Type Time to maximal ALC growth rate (days post-transplant) in patients demonstrating different recovery kinetics.

| Pattern | N | $a$<br>(Mean) | SE | 95% CI |
| --- | --- | --- | --- | --- |
| A | 65 | 22.4 | 1.56 | 19.3, 25.5 |
| B | 50 | 22.0 | 1.80 | 18.5, 25.6 |
| C | 16 | 16.9 | 3.63 | 9.7, 24.0 |

**TABLE S3.** Logistic parameter R and Pattern Type. Lymphoid growth rates in patients demonstrating different ALC recovery kinetics.

| Pattern | N | (R)<br>Mean | SE | 95% CI | p-value |
| --- | --- | --- | --- | --- | --- |
| A | N | Mean | SE | 95% CI | 0.14 |
| B | 65 | 0.26 | 0.114 | 0.03, 0.48 |  |
| C | 50 | 0.53 | 0.132 | 0.27, 0.79 |  |

**Figure S6:**
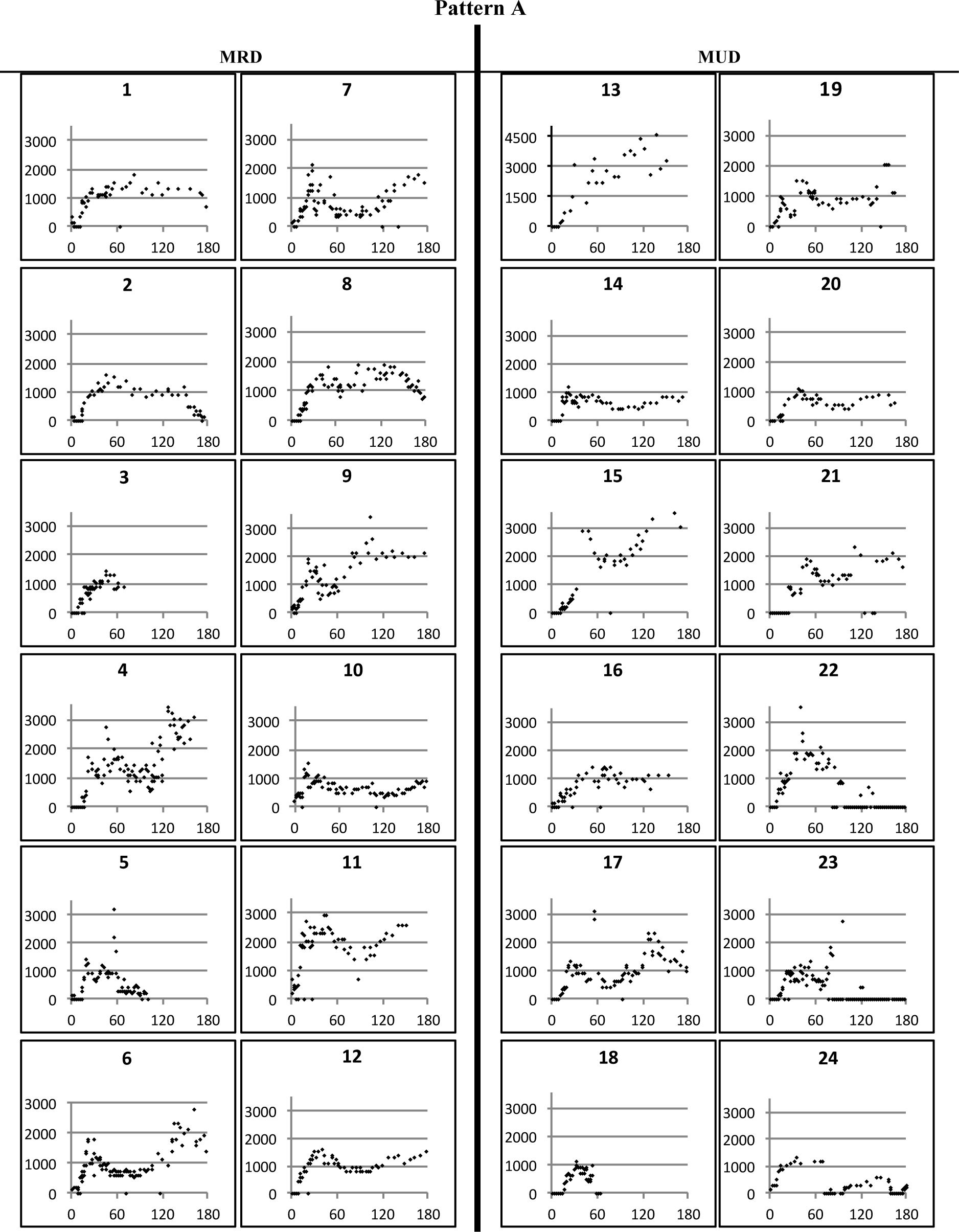
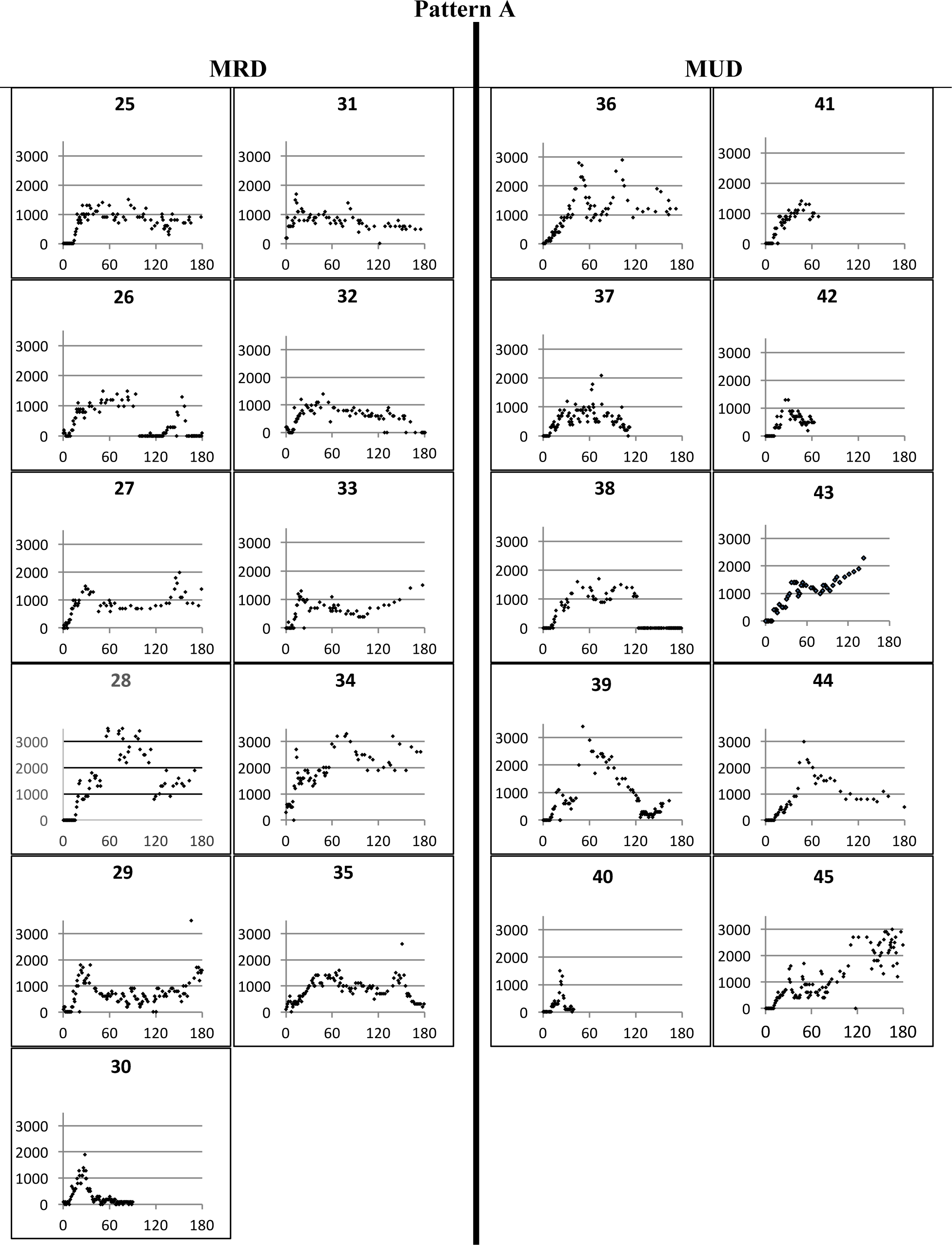
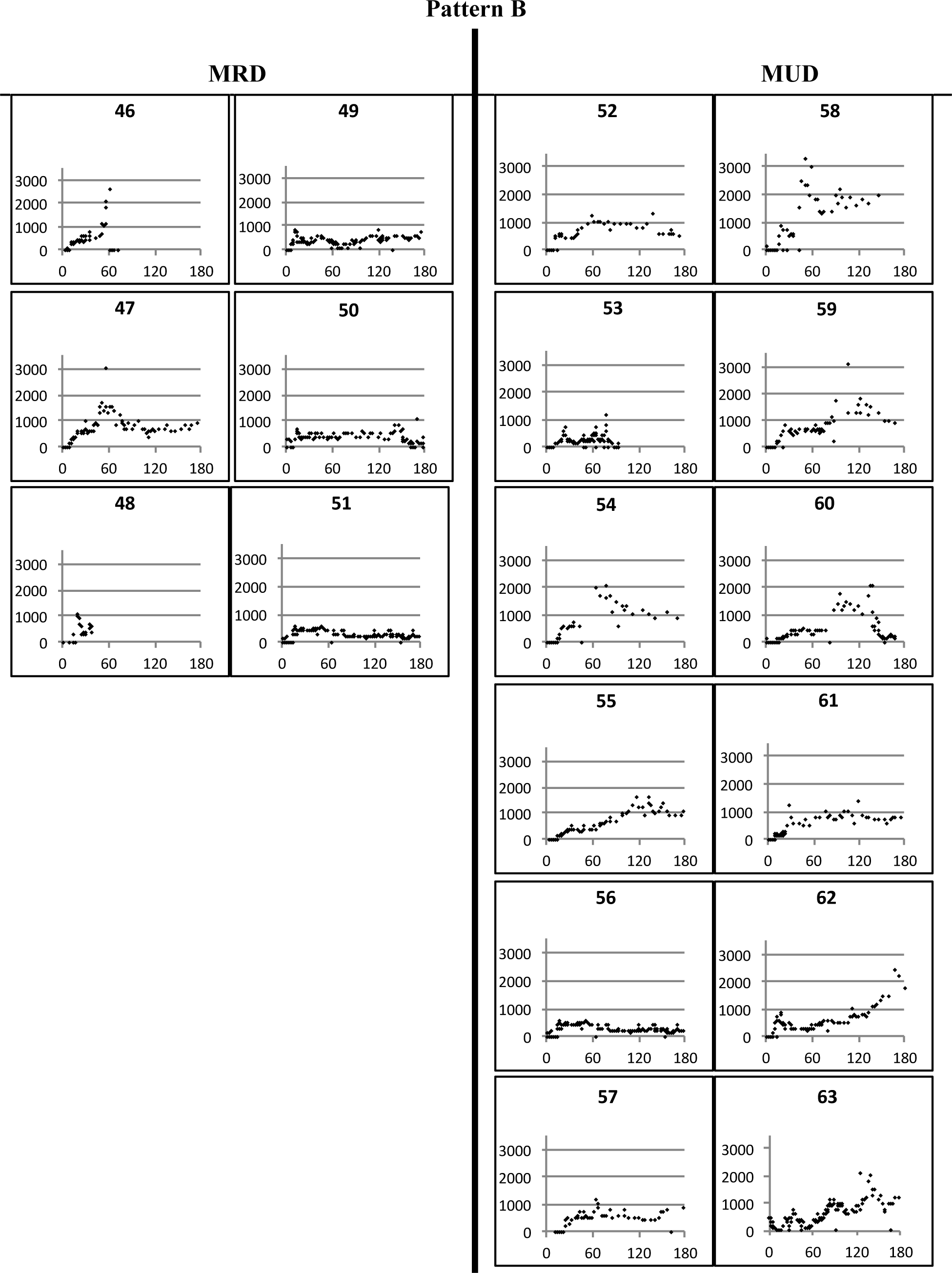
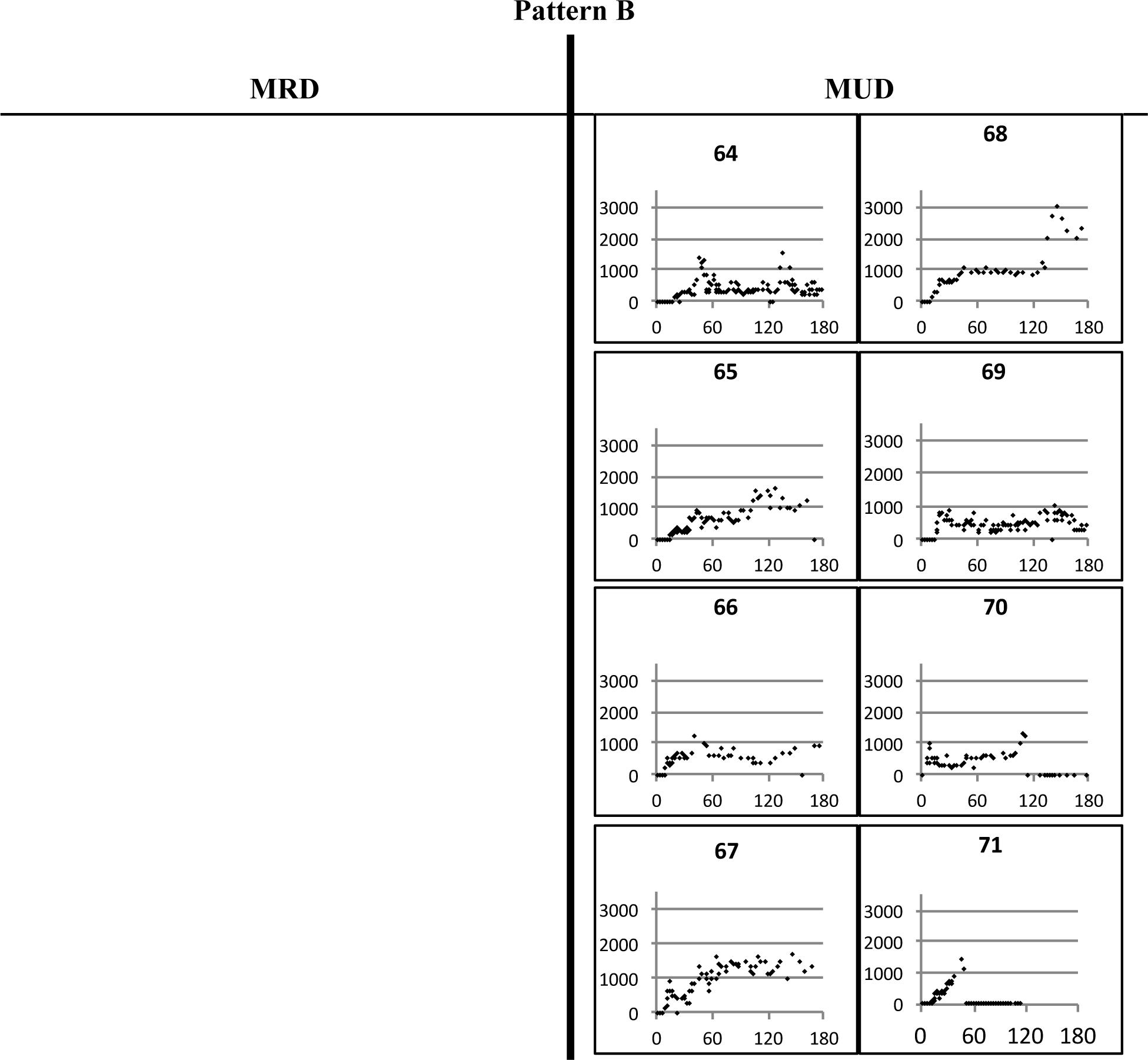
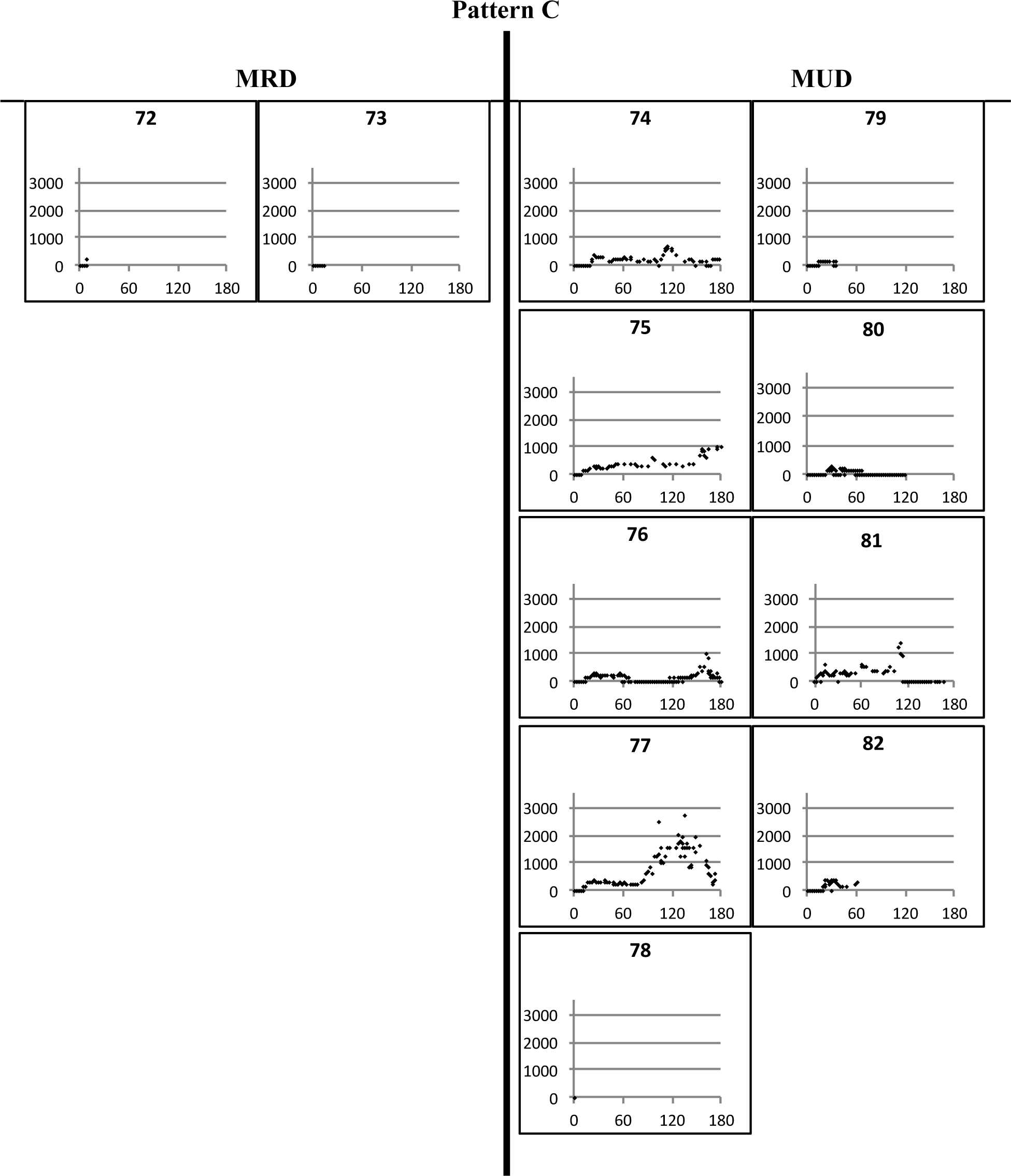
Lymphoid Reconstitution. Graphed are ALC (y-axis) and days post transplant (x-axis) in all patients

## REFERENCES

1 BirnbaumA.On the foundations of statistical inference. Journal of the American Statistical Association, 1962, 298(57) 269.

2 Biggs, N. Discrete Mathematics(2012). Oxford University Press, London, UK.

3 LaurinD, HannaniD, PernolletM, MoineA, PlumasJ, BensaJC, CahnJY, GarbanF.Immunomonitoring of graft---versus---host minor histocompatibility antigen correlates With graft---versus---host disease and absence of relapse after graft. Transfusion. 2010Feb;50(2): 418–28.

4 Morrrison F. The Art of Modeling Dynamic Systems:Forecasting for Chaos, Randomness and Determinism. Stochastic systems---Pattern IVp. 278. 2008, Dover publications, Mineola, NY.

5 ToorAA, KobulnickyJD, SalmanS, RobertsCH, Jameson---LeeM,MeierMeier J, ScaloraA,ShethN, KopardeV, SerranoM,BuckGA, ClarkWB, McCartyJM, ChungHM, ManjiliMH, Sabo RT and Neale MC. Stem Cell transplantation As a dynamical system: are clinical outcomes deterministic? Frontiers in Immunology, 2014; 5:613. doi: 10.3389/fimmu.2014.00613

6 BarrettJ.Improving outcome of allogeneic stem cell transplantation by immunomodulation Of the early post---transplant environment. Curr Opin Immunol(2006) 18:592–8.

7 Vandermeer, J.H.; Goldberg, D. E.(2003). Population ecology: First principles. Woodstock, Oxfordshire: Princeton University Press.ISBN 0–691–11440–4.

8 Toor,AmirA. et al.“Stem Cell Transplantation as a Dynamical System: Are Clinical Outcomes Deterministic?”Frontiers in Immunology 5(2014): 613.PMC. Web. 11July2016.

9 I. Stewart The imbalance of nature chaos theory. In Pursuit of the Unknown, 17 Equations that Changed the World.Philadelphia, PA: Basic Books; (2012). 283p

10 MayRM.. Biological populations with nonoverlapping generations: stable points, stable cycles, and chaos. Science(1974) 186: 645–647. 10.1126/science.186.4164.645

11 Savani BN,Mielke S, Rezvani K,Montero A, YongAS, WishL, et al.Absolute lymphocyte Count on day 30 is a surrogate for robust hematopoietic recovery and strongly predicts Outcome after T cell---depleted allogeneic stem cell transplantation. Biol Blood Marrow Transplant(2007) 13:1216–23.

12 KawaseT, MorishimaY, MatsuoK, KashiwaseK,InokoH, SajiHet al.High-risk HLA allele mismatch combinations responsible for severe acute graft-versus-host disease and implication for its molecular mechanism. Blood2007; 110: 2235–2241.

13 LeeSJ, KleinJ, HaagensonM,Baxter-LoweLA,ConferDL, EapenM et al.Highresolution donor-recipient HLA matching contributes to the success of unrelated donor marrow transplantation. Blood2007; 110: 4576–4583.

14 KawaseT,MatsuoK, KashiwaseK,InokoH,SajiH, OgawaSet al.HLA mismatch combinations associated with decreased risk of relapse: implications for the molecular mechanism. Blood2009; 113: 2851–2858

15 PericZ, CahuX, ChevallierP, BrissotE, MalardF, GuillaumeTet al.Features of Epstein-Barr Virus (EBV) reactivation after reduced intensity conditioning allogeneic hematopoietic stem cell transplantation. Leukemia2011; 25: 932–938.

16 DeegH, StorerB, BoeckhM, MartinP, McCuneJ, MyersonDet al.Reduced incidence of acute and chronic graft-versus-host disease with the addition of thymoglobulin to a targeted busulfan/cyclophosphamide regimen. Biol Blood Marrow Transplant2006; 12: 573–584.

17 DugganP, BoothK, ChaudhryA, StewartD, RuetherJ, GluckS et al.Unrelated donor BMT recipients given pretransplant low-dose antithymocyte globulin have outcomes equivalent to matched sibling BMT: a matched pair analysis. Bone Marrow Transplant2002; 30: 681–686.

18 DeegH, StorerB, BoeckhM, MartinP, McCuneJ, MyersonDet al.Reduced incidence of acute and chronic graft-versus-host disease with the addition of thymoglobulin to a targeted busulfan/cyclophosphamide regimen. Biol Blood Marrow Transplant2006; 12: 573–584.

19 BacigalupoA, LamparelliT, BarisioneG, BruzziP, GuidiS,AlessandrinoPEet al.Thymoglobulin prevents chronic graft-versus-host disease, chronic lung dysfunction, and late transplant-related mortality: long-term follow-up of a randomized trial in patients undergoing unrelated donor transplantation. Biol Blood Marrow Transplant2006; 12: 560–565.

20 PortierDA, SaboRT, RobertsCH, FletcherDS, MeierJ, ClarkWB,NealeMC,ManjiliMH,McCartyJM, ChungHM,ToorAA.Bone Marrow Transplant. 2012Dec;47(12): 1513–9.doi: 10.1038/bmt.2012.81. Epub2012May14.

21 BacigalupoA, LamparelliT, BruzziP, GuidiS, AlessandrinoPE, diBartolomeo Pet al.Antithymocyte globulin for graft-versus-host disease prophylaxis in transplants from unrelated donors: 2randomized studies from Gruppo Italiano Trapianti Midollo Osseo (GITMO). Blood2001; 98: 2942–2947.

22 BacigalupoA, BallenK, RizzoD, et al.Defining the intensity of conditioning regimens: working definitions. Biol Blood Marrow Transplant. 2009;15(12): 1628–1633.

23 RowlingsPA, PrzepiorkaD, KleinJP, et al.IBMTR Severity Index for grading acute graft-versus-host disease: retrospective comparison with Glucksberg grade. Br J Haematol1997; 97:855.

24 RowlingsPA, PrzepiorkaD, KleinJP, et al.IBMTR Severity Index for grading acute graft-versus-host disease: retrospective comparison with Glucksberg grade. Br J Haematol1997; 97:855.

25 ShulmanHM,SullivanKM, WeidenPL, et al.Chronic graft-versus-host syndrome in man. A long-term clinicopathologic study of 20 Seattle patients. Am J Med1980; 69:204.

26 KimH, ArmandP, FrederickD, Andler E, Cutler C, Koreth J,AlyeaE,AntinJ, Soiffer J, Ritz J Ho V. Lymphocyte Count Recovery after Allogeneic Hematopoietic Stem Cell Transplantation Predicts Clinical Outcome. Biology of Blood and Marrow Transplantation, May2015, Vol.21(5), pp.873–880

27 KimHT, FrederickD, AndlerE, et al.Prognostic signifcance of whiteblood cell counts on clinical outcome after allogeneic hematopoietic cell transplantation. Am J Hematol. 2014;89:591–597

28 YamamotoW, OgusaE, MatsumotoK, et al.Lymphocyte recovery onday 100 after allogeneic hematopoietic stem cell transplant predicts non-relapse mortality in patients with acute leukemia or myelodys-plastic syndrome. Leuk Lymphoma. 2014;55:1113–1118.

